# Human H-ferritin presenting RBM of spike glycoprotein as potential vaccine of SARS-CoV-2

**DOI:** 10.1101/2020.05.25.115618

**Authors:** Dehui Yao, Fang Lao, Zeyi Zhang, Yan Liu, Jianwei Cheng, Fengjiao Ding, Xiaofei Wang, Lun Xi, Chuang Wang, Xichong Yan, Rongkun Zhang, Fangxing Ouyang, Hui Ding, Tianyi Ke

**Author notes:** ^*^Corresponding authors: Tianyi Ke; Hui Ding.

## Abstract

The outbreak of COVID-19 has so far inflicted millions of people all around the world and will have a long lasting effect on every aspect of everyone’s life. Yet there is no effective approved treatment for the disease. In an effort of utilizing human ferritin as nanoplatform for drug delivery, we engineered a fusion protein by presenting receptor-binding motif (RBM) of SARS-CoV-2 virus spike glycoprotein on the N-terminus of ferritin subunits. The designed fusion protein with a cage-like structure, similar to that of corona virus, is a potential anti-SARS-CoV-2 vaccine. We hereby show the construction, preparation, and characterization of the fusion protein RBM-HFtn. Our initial affinity study confirmed its biological activity towards ACE2 receptor which suggests its mode of action against SARS-CoV-2 could be either through vaccine therapy or blocking the cellular entry of virus as antagonist of ACE2 receptor.

## Introduction

The pandemic coronavirus disease (COVID-19) by SARS-CoV-2 virus has caused tremendous suffering to tens of millions of people around the world. Even though quite a few clinical studies involving different approaches are undergoing for the treatment of the disease, there is no effective cure yet up to date. Vaccines therefore is urgently needed for the preventing further spread of the COVID-19.

The corona virus, SARS-CoV-2, consists of a large RNA genome, four structural proteins, 16 nonstructural proteins, and some accessory proteins. The four structural proteins include spike, envelope, membrane, and nucleocapsid proteins, of which the spike glycoprotein is of particular interest for it is a popular vaccine target for corona virus. Antibodies targeting the spike glycoprotein of SARS-CoV and MERS-CoV, especially its receptor-binding domain (RBD), was found to efficiently neutralize virus infection [1, 2]. Antibodies from SARS-CoV and SARS-CoV-2 patients however showed limited cross neutralization, in spite of the high sequence similarity between two viruses [3]. Other vaccine approaches include the production of live attenuated whole virion vaccines, inactivated whole virion vaccines, recombinant protein vaccines, and mRNA based vaccines [4].

SARS-CoV-2 was found to enter cells through binding of the host cellular receptor angiotensin-converting enzyme 2 (ACE2) via its spike glycoprotein soon after its outbreak in China [3]. Later the cryo-EM revealed the structure of SARS-CoV-2 S-RBD complexed with its receptor human ACE2 [5–8]. The S1 subunit of spike glycoprotein undergoes a hinge-like conformational transition from “down” conformation to “up” conformation before binding to ACE2 [8]. The receptor-binding motif (RBM, 72 amino acid in total) of spike glycoprotein in close contact of ACE2 receptor was identified for its sequence between 437 – 508 [9], and this RBM was consistent with the identified S-RBD [8]. This conformational transition state has become the target for antibody-mediated neutralization, and the atomic-level understanding of this transitional state and the identification of S-RBM would facilitate the vaccine design and development against SARS-CoV-2.

Ferritin is a 24-mer protein assembly consisting of heavy chain (21 kD) and light chain (19 kD). It has a cage-like structure in a way similar to SARS-CoV-2. Because of its unique structure, ferritin is a promising nanoplatform for antigen presentation and immune stimulation [10–13]. Its spherical architecture has an outer diameter of 12 nm, suitable for rapid tissue penetration and draining to lymph node [14]. In this work, we engineered a human ferritin heavy chain (HFtn) by fusing and presenting the RBM of its spike glycoprotein as potential vaccine of SARS-CoV-2.

## Materials and methods

### Materials

Human ACE2 (his tag) (10108-H08H), ACE2 rabbit Antibody (10108-RP01), goat anti-Rabbit/HRP secondary antibody (SSA004, and SARS-CoV-2 (2019-nCoV) and RBD of spike glycoprotein (mFC tag)(40592-V05H) were purchased from Sino Biological (Beijing, China).

### Construction of the gene of RBM-HFtn fusion protein

The receptor binding motif (RBM) of SARS-CoV-2 spike glycoprotein (72 amino acids, sequence from 439 – 508) was incorporated into the N-terminus of human H-ferritin using a linker of (GGGGS)_3_ to form a RBM-HFtn fusion protein. The sequence of RBM peptide, RBM-HFtn fusion protein, and the DNA sequence (code: XYD-403-000) for optimized expression in *E. Coli* is described in Supplementary Methods. The gene synthesis and plasmid construction was completed by Shanghai Generay Biotech Co., Ltd (Shanghai, China) using *p*ET-22b(+) plasmid vector with Amp^+^ resistance and two enzyme cleavage sites Nde I and Bam H I (Supplementary Figure S1). The plasmid construction was confirmed by enzyme cleavage mapping and gene sequencing.

### Cloning, expression, and refolding of fusion protein RBM-HFtn

*E.coli* BL21 (DE3) was transformed with *p*ET-22b(+) plasmid containing XYD-403-000 gene, Ampicillin positive single colonies were picked and cultured in LB broth at 37°C overnight. Then, RBM-HFtn fusion protein production was induced by 0.5 mM isopropyl-β-D-thiogalactoside (IPTG, Inalco, USA), and cells were incubated for an additional 5 h at 25°C. After incubation, the *E.coli* cells were harvested at 10,000 rpm at 4°C for 30 min and the pellets were resuspended in Tris buffer (20 mM Tris, pH 8.0) and the resuspended *E.coli* cells was ruptured at 1000 bar for three times and centrifuged at 10,000 rpm at 4°C for 30 min. The target protein (molecular weight 30.4 kD) was found to be in precipitates, suggesting it exists mainly in the form of inclusion bodies. Inclusion bodies of RBM-HFtn subunits were refolded by first dissolved in 8M urea followed by gradual dialysis with 6, 4, 2, 1, 0.5, 0 M urea and 20 mM Tris-HCl buffer with 500 mM NaCl, pH 8.0. The refolded protein solution was centrifuged at 5,000 g at 4°C for 30 min and supernatant was collected.

### Purification of fusion proteins

The refolding solution was concentrated and buffer exchanged to 25 mM Tris-HCl buffer (pH 8.0) with a Millipore lab scale tangential flow filter (TFF) system. After buffer exchange, RBM-HFtn was found to have a typical soluble ferritin structure. The fusion protein with its right space structure was further purified by using anion exchange chromatography (flow through mode), followed with a size-exclusion chromatography to remove aggregates and other low molecule impurities.

### Characterization of fusion protein RBM-HFtn

The physico-chemical properties of fusion RBM-HFtn were analyzed by SEC-HPLC (Agilent 1260 Infinity II HPLC system, column Agilent AdvancedBio SEC 300Å 2.7 μm 7.8 × 300mm) with mobile phase 50 mM Tris buffer (pH 8.0) at flow rate 0.5 mL/min and detection wavelength 280 nm. Hydrodynamic size analysis by Nano ZSE Nanosizer (Malvern, UK) using automatic mode with material as protein and dispersant as 50 mM Tris buffer (pH 8.0). The morphology and size of fusion protein was further analyzed by transmission electron microscopy (TEM) (FEI Tecnai Spirit (100KV)) after staining with 2% uranyl acetate.

### Binding affinity analysis by ELISA

The binding affinity of RBM-HFtn towards ACE2 receptor was analyzed with indirect ELISA. Briefly, varied concentrations of RBM-HFtn fusion protein and S-RBD (positive control) in 100 μL coating buffer were coated onto MaxiSorp plate (CORN) by incubation at 4°C overnight. After blocking with 5% BSA, ACE2 protein (concentration 0.5 μg/ml) was allowed to bind to the surface coated proteins for 2 h at 37°C, followed by washing with washing buffer three times. Anti-ACE2 antibody (dilution 1:1000) was incubated for 1.5 hours at 37°C followed by washing for three times. Secondary antibody/HRP (1:5000 dilution) in 5% BSA was incubated 37°C for 0.5 h followed by washing for three times. The TMB substrate was added and incubated for 30 min, and then the absorbance was read at 650 nm wavelength.

## Results and discussion

### Construction of RBM-HFtn

Ferritin heavy chain protein has found many biological applications in nanomedicines and molecular diagnostics [15–17]. Lately, a few constructs based on ferritin have been developed as antivirus and anticancer vaccines [10, 11, 13]. Because ferritin heavy chain protein is derived from a natural occurring protein in human, itself has low immunogenicity. Its 24-mer assembly cage-like structure has four different symmetries, six 4-fold axes, eight 3-fold axes, twelve 2-fold axes, and twenty-four C3-C4 interfaces. By presenting a fusion protein at the 3-fold axes, ferritin was able to display eight trimers of fused protein, resulting in enhanced immunogenicity of protein on display [18]. In this work we engineered a human ferritin heavy chain fused with the RBM of spike glycoprotein of SARS-CoV-2 at its N-terminus with (GGGGS)_3_ short peptide linker (Figure 1A). We chose a short polypeptide sequence (72 amino acid) of spike glycoprotein RBM (S-RBM) based on latest 3-D cryo-EM studies of spike glycoprotein-ACE2 complex [8, 9]. Because this sequence doesn’t appear to have a stable 3-D structure, we engineered it at the N-terminus, so that the fused S-RBM subunits are distant from each other and won’t affect the formation 24-mer assembly. For the same reason, we chose a minimal size of RBD so that it is properly displayed on the surface of ferritin. In spite of those considerations, the RBM-HFtn fusion protein was found mainly in the inclusion bodies, instead of in soluble form.

**Figure 1.**
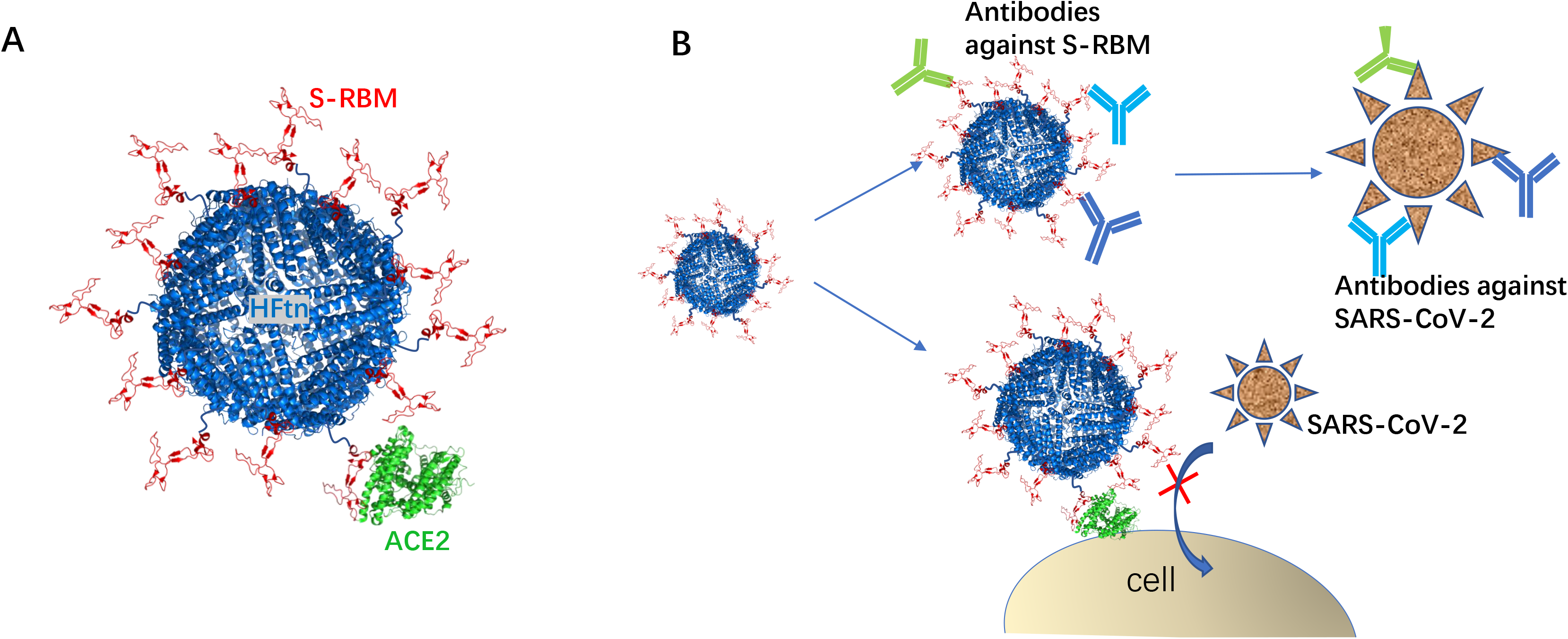
Schematic representation of RBM-HFtn structure with ferritin as scaffold (blue) fused with RBM of SARS-CoV-2 spike glycoprotein (red); The RBM-HFtn is capable of complexing with ACE2 (green) (A). Total 24 copies of RBM are presented on the ferritin surface, here only 14 copies were shown for symbolic representation. Two hypothetical antivirus pathways of RBM-HFtn by inducing antibodies against SARS-CoV-2 (B, upper pathway) and/or by blocking virus entry through competitive binding to ACE2 (B, lower pathway).

Hypothetically, the designed RBM-HFtn fusion protein may involve in two different pathways against virus infection. First and most importantly, the RBM-HFtn may act as vaccine against SARS-CoV-2, and antibodies responsive to spike glycoprotein RBM may subsequently neutralize SARS-CoV-2, followed by removal of virus by immune system (Figure 1B, upper pathway). Two antibodies found in COVID-19 patients were shown to block the binding between virus S-protein RBD and cellular receptor ACE2 [19]. Their therapeutic effect was validated in mouse model by reducing virus titer in infected lungs [19]. Those findings supported our hypothesis that by properly presenting S-protein RBD/RBM, the designed RBM-HFtn has great potential as an anti-SARS-CoV-2 vaccine, thus inducing the production of antivirus antibodies. The second pathway, more obvious and direct, is to block the cellular entry of SARS-CoV-2 by preoccupy the ACE2 receptor with RBM-HFtn and suppress the virus proliferation (Figure 1B, lower pathway). The first pathway is more effective for prevention of virus infection. The second pathway is valuable for treatment after virus infection as antagonist of ACE2 receptor.

### Preparation, purification and characterization of RBM-HFtn

The expressed RBM-HFtn was found in the form of inclusion bodies in the bacteria lysis precipitates (Supplementary Figure S2) with apparent molecular weight of 30 kD which is consistent with the size of RBM-HFtn subunits. The protein was properly refolded and reassembled to 24-mer protein by gradual dialysis in urea buffers of decreased concentration, and finally in 20 mM Tris-HCl buffer with 500 mM NaCl, pH 8.0. The refolded protein was sequentially purified with a Millipore lab scale tangential flow filter (TFF) system, anion exchange chromatography (flow through mode), and a size-exclusion chromatography to remove host cell protein, host cell DNA, endotoxin, protein aggregates and other low molecule impurities. The purified RBM-HFtn was found have a high purity 98% as evaluated by size exclusion chromatography (SEC) (Figure 2.) It has slightly increased hydrodynamic size of 15.0 ± 0.6 nm measured by Nanosizer in comparison with the known size of HFtn,12 nm (Figure 2B). Transmission electron microscopy confirmed the size of RBM-HFtn fusion protein in the range of 14-16 nm with almost uniform morphology of round shape (Figure 2C). The SEC, Nanosizer, and TEM results confirmed the successful construct of the RBM-HFtn.

**Figure 2.**
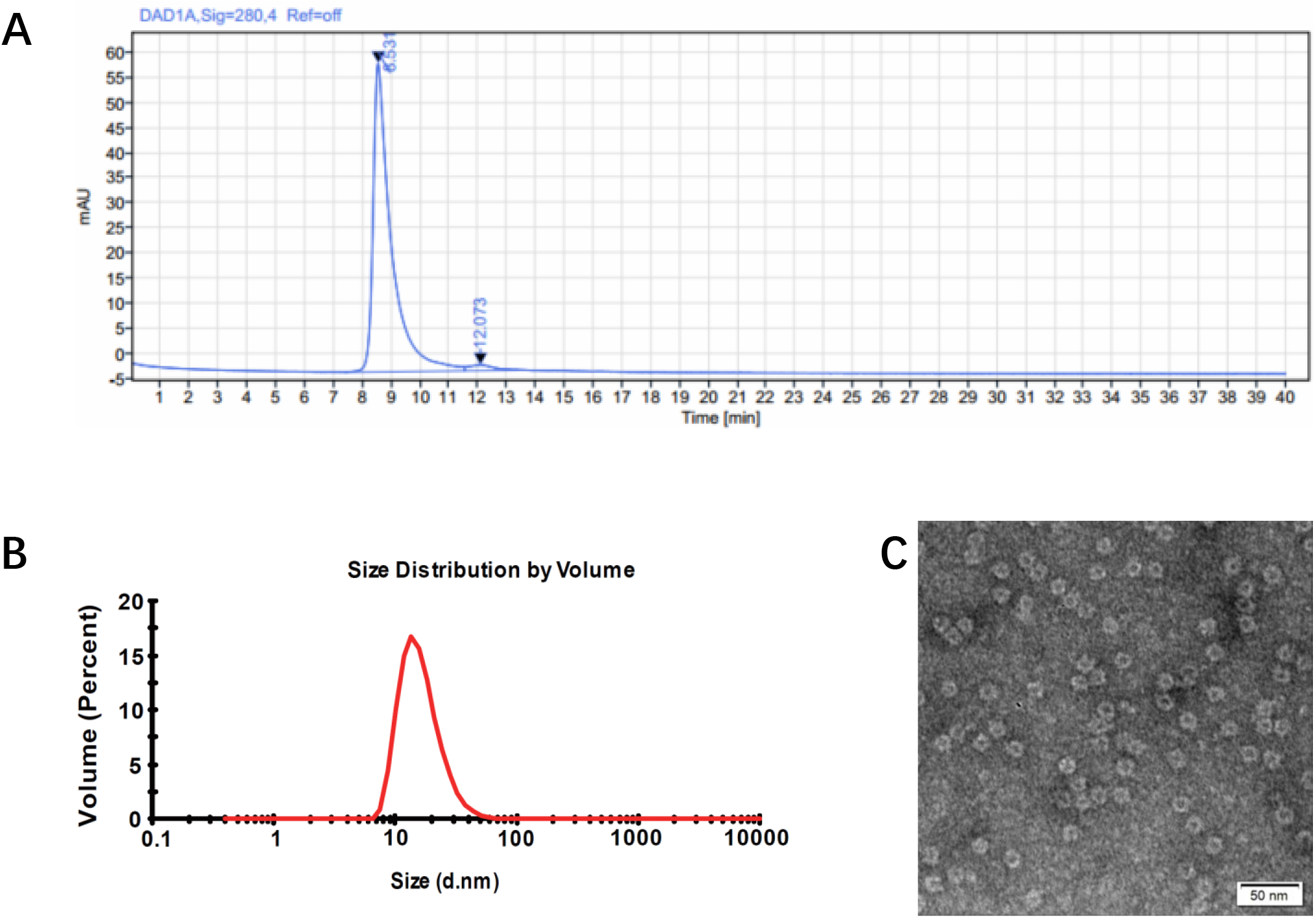
Characterization of RBM-HFtn. SEC of RBM-HFtn detected at 280 nm showing its purity of 98% (A). Hydrodynamic diameter of RBM-HFtn measured by Nanosizer as 15.0 ± 0.6 nm (B). Morphology and size determination by TEM (C).

### Binding affinity towards ACE2 by indirect ELISA

RBD of Spike glycoprotein (mFc tag) and RBM-HFtn of varied concentration in 100 μL was incubated in MaxiSorp plate and ACE2 was allowed to bind to the surface coated proteins. Their binding was detected by ACE2 antibody followed by its secondary antibody. In this indirect ELISA, the maximum binding intensity correlates with the binding site on surface. As indicated by the results in Figure 3, S-RBD showed higher plateau than RBM-HFtn, suggesting more binding sites are available in S-RBD coated wells than in RBM-HFtn coated wells. The EC50s of the binding between ACE2 and S-RBD, RBM-HFtn were estimated to be 60.66 nM and 23.36 nM respectively. Apparently RBM-HFtn has higher binding affinity than S-protein in binding to their same ACE2 receptor. The apparent higher binding affinity may attribute to the cage-like structure of RBM-HFtn, which presents multiple copies (24 copies) of RBM on the surface. This result suggested that the RBM is properly presented on surface of heavy chain human ferritin and is recognizable by the ACE2 receptor. Its potential as SARS-CoV-2 vaccine and as antagonist of ACE2 receptor is being further studied in animal experiments.

**Figure 3.**
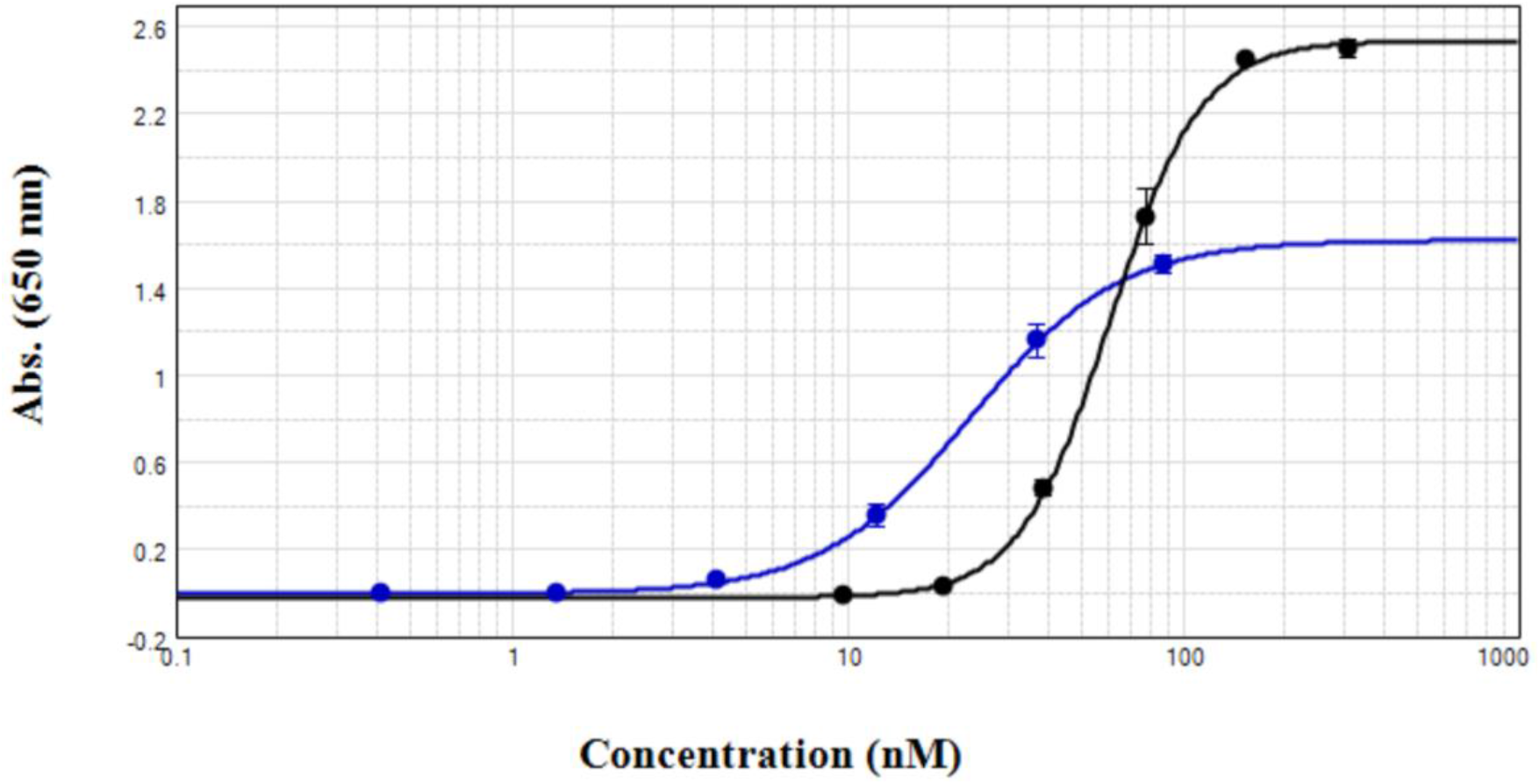
Analysis of ACE2 binding to surface coated RBD of SARS-CoV-2 (2019-nCoV) Spike glycoprotein (mFC tag) (black dots) and RBM-HFtn (blue square) of varied concentrations by ELISA.

## Conclusions

The receptor-binding motif (RBM) of SARS-CoV-2, a 72-amino acid polypeptide, was fused with N-terminus of human heavy chain ferritin (HFtn) through a proper linker. The constructed RBM-HFtn was found in inclusion bodies of bacterial lysis and was able to refold by gradual dialysis. The purified RBM-HFtn was found to be in good purity, expected size and morphology. Its biological activity towards ACE2 receptor was confirmed by ELISA and the fusion protein RBM-HFtn bodes well as potential vaccine and therapeutics against SARS-CoV-2 as ACE2 receptor antagonist.

## Authors’ contributions

D. Yao, Z. Zhang, F. Ding, X. Wang, L. Xi: construction of the gene and production of fusion protein

F. Lao, T. Ke: design and supervision of the experiment

Y. Liu, C. Wang: biological evaluation of fusion protein

H. Ding: writing and revision of the manuscript and data analysis

J. Cheng, R. Zhang, X. Yao: purification of fusion protein

F. Ouyang: administrative and material support

## References

1. Du, L., et al., The spike protein of SARS-CoV--a target for vaccine and therapeutic development. Nat Rev Microbiol, 2009. 7(3): p. 226–36.

2. Wang, Q., et al., MERS-CoV spike protein: Targets for vaccines and therapeutics. Antiviral Res, 2016. 133: p. 165–77.

3. Zhou, P., et al., A pneumonia outbreak associated with a new coronavirus of probable bat origin. Nature, 2020. 579(7798): p. 270–273.

4. Amanat, F. and F. Krammer, SARS-CoV-2 Vaccines: Status Report. Immunity, 2020. 52(4): p. 583–589.

5. Lan, J., et al., Structure of the SARS-CoV-2 spike receptor-binding domain bound to the ACE2 receptor. Nature, 2020.

6. Yan, R., et al., Structural basis for the recognition of SARS-CoV-2 by full-length human ACE2. Science, 2020. 367(6485): p. 1444–1448.

7. Shang, J., et al., Structural basis of receptor recognition by SARS-CoV-2. Nature, 2020.

8. Wrapp, D., et al., Cryo-EM structure of the 2019-nCoV spike in the prefusion conformation. Science, 2020. 367(6483): p. 1260–1263.

9. Wan, Y., et al., Receptor Recognition by the Novel Coronavirus from Wuhan: an Analysis Based on Decade-Long Structural Studies of SARS Coronavirus. J Virol, 2020. 94(7).

10. Kanekiyo, M., et al., Self-assembling influenza nanoparticle vaccines elicit broadly neutralizing H1N1 antibodies. Nature, 2013. 499(7456): p. 102–6.

11. Lee, B.R., et al., Engineered Human Ferritin Nanoparticles for Direct Delivery of Tumor Antigens to Lymph Node and Cancer Immunotherapy. Sci Rep, 2016. 6: p. 35182.

12. Li, Z., et al., A milk-based self-assemble rotavirus VP6-ferritin nanoparticle vaccine elicited protection against the viral infection. J Nanobiotechnology, 2019. 17(1): p. 13.

13. Wang, L., et al., Structure-based design of ferritin nanoparticle immunogens displaying antigenic loops of Neisseria gonorrhoeae. FEBS Open Bio, 2017. 7(8): p. 1196–1207.

14. Bachmann, M.F. and G.T. Jennings, Vaccine delivery: a matter of size, geometry, kinetics and molecular patterns. Nat Rev Immunol, 2010. 10(11): p. 787–96.

15. Truffi, M., et al., Ferritin nanocages: a biological platform for drug delivery, imaging and theranostics in cancer. Pharmacological Research, 2016.

16. Zhen, Z., et al., Ferritins as nanoplatforms for imaging and drug delivery. Expert Opinion on Drug Delivery, 2014. 11(12): p. 1913–1922.

17. Chiou, B. and J.R. Connor, Emerging and Dynamic Biomedical Uses of Ferritin. Pharmaceuticals (Basel), 2018. 11(4).

18. Sliepen, K., et al., Structure and immunogenicity of a stabilized HIV-1 envelope trimer based on a group-M consensus sequence. Nat Commun, 2019. 10(1): p. 2355.

19. Wu, Y., et al., A noncompeting pair of human neutralizing antibodies block COVID-19 virus binding to its receptor ACE2. Science, 2020.

