## Supplementary for "Human H-ferritin presenting RBM of spike glycoprotein as potential vaccine of SARS-CoV-2"

**Methods**

***Construction of RBM-HFn fusion protein***

RBM of SARS-CoV-2 spike glycoprotein sequence

NSNNLDSKVGGNYNYLYRLFRKSNLKPFERDISTEIYQAGSTPCNGVEGFNCYFPLQSY
GFQPTNGVGYQPY

RBM-HFtn fusion protein sequence.

MNSNNLDSKVGGNYNYLYRLFRKSNLKPFERDISTEIYQAGSTPCNGVEGFNCYFPLQS
YGFQPTNGVGYQPYGGGGSGGGGSGGGGSTTASTSQVRQNYHQDSEAAINRQINLEL
YASYVYLSMSYYFDRDDVALKNFAKYFLHQSHEEREHAEKLMKLQNQRGGRIFLQDIKK
PDCDDWESGLNAMECALHLEKNVNQSLLELHKLATDKNDPHLCDFIETHYLNEQVKAIK
ELGDHVTNLRKMGAPESGLAEYLFDKHTLGDSDNES

Note: yellow shaded sequence is the linker between the HFtn and RBM

Gene sequence of RBM-HFtn fusion protein (gene code XYD-403-000).

ATGAATAGTAATAATCTGGATTCTAAAGTGGGCGGCAATTATAATTATCTGTATCGCCTG
TTTCGTAAATCAAATCTGAAACCGTTTGAACGCGATATTAGTACCGAAATTTATCAGGC
AGGCTCTACCCCGTGTAATGGTGTTGAAGGCTTTAATTGTTATTTTCCGCTTCAAAGC
TATGGCTTTCAGCCGACCAATGGCGTTGGCTATCAGCCGTATGGTGGTGGCGGTTCA
GGCGGCGGTGGTAGCGGCGGTGGCGGTAGTACCACCGCAAGCACCTCACAGGTTC
GTCAGAATTATCATCAGGATAGCGAAGCAGCAATTAATCGCCAGATTAATCTGGAACT
GTATGCAAGCTATGTGTATCTGAGTATGTCTTATTATTTTGATCGCGATGATGTTGCACT
GAAAAATTTTGCAAAATATTTTCTGCATCAGTCTCATGAAGAACGCGAACATGCAGAA
AAACTGATGAAACTCCAAAATCAGCGTGGTGGTCGCATTTTTCTTCAAGATATTAAAAA
ACCGGATTGTGATGATTGGGAAAGTGGCCTGAATGCAATGGAATGTGCACTGCATCT
GGAAAAAAATGTTAATCAGTCACTGCTGGAACTGCATAAACTGGCAACCGATAAAAAT
GATCCGCATCTGTGTGATTTTATTGAAACCCATTATCTGAATGAACAGGTTAAAGCAAT
TAAAGAACTGGGTGATCATGTGACCAATCTGCGTAAAATGGGCGCACCGGAAAGCG
GCCTGGCAGAATATCTGTTTGATAAACATACCCTGGGCGATAGCGATAATGAAAGT

### Figures

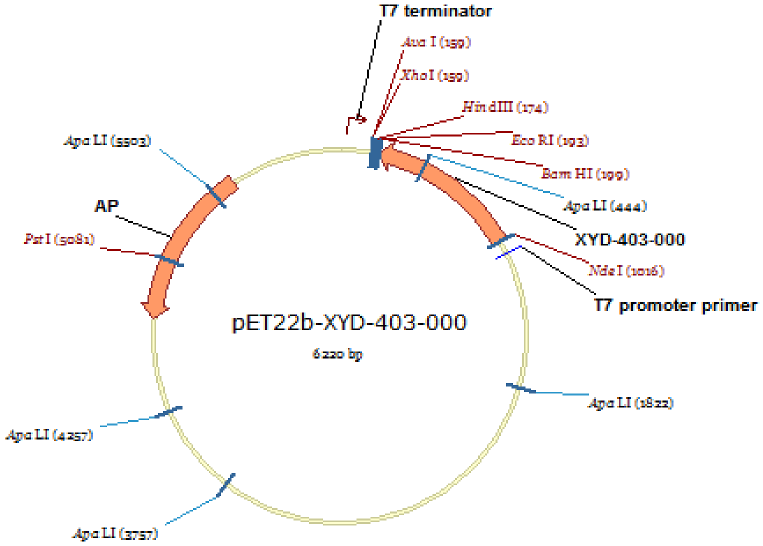


Figure S1. Construction of *p*ET-22b(+) plasmid containing XYD-403-000 gene.

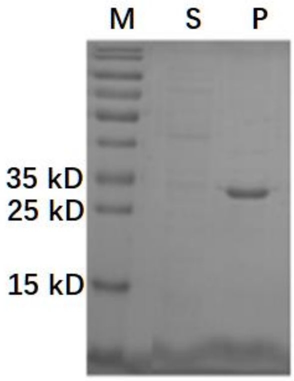


Figure S2. RBM-HFtn expression results by denatured SDS-PAGE. Lane M is protein ladder; Lane S is bacteria lysis supernatant; Lane P is bacteria lysis precipitates.
